# Dose optimization of an adjuvanted peptide-based personalized neoantigen melanoma vaccine

**DOI:** 10.1101/2023.06.09.544293

**Authors:** Wencel Valega-Mackenzie, Marisabel Rodriguez Messan, Osman N. Yogurtcu, Ujwani Nukala, Zuben E. Sauna, Hong Yang

## Abstract

The advancements in next-generation sequencing have made it possible to effectively detect somatic mutations, which has led to the development of personalized neoantigen cancer vaccines that are tailored to the unique variants found in a patient’s cancer. These vaccines can provide significant clinical benefit by leveraging the patient’s immune response to eliminate malignant cells. However, determining the optimal vaccine dose for each patient is a challenge due to the heterogeneity of tumors. To address this challenge, we formulated a mathematical dose optimization problem that aims to find the optimal personalized vaccine doses for a given fixed vaccination schedule, based on a previous mathematical model that encompasses the immune response cascade produced by the vaccine in a patient. To validate our approach, we performed *in silico* experiments on six patients with advanced melanoma. We compared the results of applying an optimal vaccine dose to those of a suboptimal dose (dose used in the clinical trial and its deviations). Our simulations revealed that an optimal vaccine may lead to a reduction in tumor size for certain patients, with higher initial doses and lower final doses. Our mathematical dose optimization offers a promising approach to determining the optimal vaccine dose for each patient and improving clinical outcomes.

## Introduction

Cancer is the second-leading cause of death globally, accounting for approximately one in every six deaths in 2018 [1, 2]. Current cancer treatment including surgery, radiotherapy, chemotherapy, and immunotherapy, can improve a patient’s clinical outcome, but long-term survival is often impacted by the immunosuppressive environment that cancer patients experience [3]. Therapeutic cancer vaccines provide clinical benefits to cancer patients by eliciting an anti-tumor immune response, increasing survival and long-term remission [4–6]. However, dosing for optimal clinical outcomes is a key challenge in the use of cancer vaccine due to tumor heterogeneity and differences between patients [4, 7, 8].

In recent years, mechanistic and quantitative system pharmacology (QSP) models have proved to be useful for understanding the complex interactions among the immune system, tumors, and therapeutic interventions [9–13]. These mathematical tools allow for modeling specific cell populations such as dendritic, memory T, helper T, cytotoxic T, or natural killer cells, as well as the tumor microenvironment [14–19], and have been used to optimize multiple cancer treatments or cancer vaccine regimens [20–25].

Due to heterogeneity among the patients, dosing regimens for individual patients are difficult to test in clinical trials. An optimal cancer vaccine dose for an individual patient may be explored using compartmental models; however, not much work has been done in this area. The goal of this study is to propose a novel approach to quantitatively determine the optimal composition including peptide and adjuvant of a personalized cancer vaccine. To achieve this goal, we propose two optimization problems using the immunological model we developed previously [26]. The first optimization problem focuses on minimizing the overall number of residual tumor cells and total vaccine exposure throughout the treatment. The second problem seeks to minimize the number of activated *CD*4^+^ and *CD*8^+^ T cells while concurrently achieving the greatest tumor cell reduction. While the first problem optimizes for efficacy (reduction of tumor cells) the second problem optimizes for both efficacy and safety (an excessive immune response may pose a potential safety risk). We apply both optimization problems to six patients with advanced melanoma [27] to investigate whether these patients could have benefited from more effective personalized vaccine doses.

## Methodology The Model

We use our compartmental model published previously [26], which captures the interactions among the human immune system, tumor burden, and a personalized neoantigen peptide cancer vaccine. We refer to this model as MRM in this paper.

The MRM model is deterministic and consists of a set of nonlinear ordinary differential equations (ODEs) with non-negative initial conditions. The model describes the key events associated with an immune reaction to a cancer vaccine at the cellular and subcellular levels. The model describes an adjuvanted peptide-based cancer vaccine (see Equations (1)-(2) in S1 Appendix) as well as cell dynamics of the immune system at the molecular (see Equations (5)-(9) in S1 Appendix) and cellular level (see Equations ((3)-(4)) and ((10)-(13) in S1 Appendix). These equations are all interconnected to represent the immune response cascade elicited by a cancer vaccine. At the molecular level, the MRM model focuses on the processing and presentation of neoantigen molecules primarily by the dendritic cells (DCs), and captures the subcellular dynamics of endosomal peptides involving major histocompatibility complex (MHC) classes I and II in DCs. At the cellular level, the model presents the evolution of immature and mature DCs, naïve and activated T cells, and tumor cells throughout the course of the treatment with the cancer vaccine. The key immunological processes at the cellular level are activation of DCs by the adjuvant, activation of naïve *CD*4^+^ and *CD*8^+^ T cells by mature DCs carrying peptide-bound (i.e., p-MHC) molecules, proliferation and differentiation of T cells, and elimination of tumor cells by activated *CD*8^+^ T cells. A flow diagram of MRM model is shown in Fig 1. A summary of all model variables with their corresponding definitions and units are shown in Table 1.

**Fig 1.**
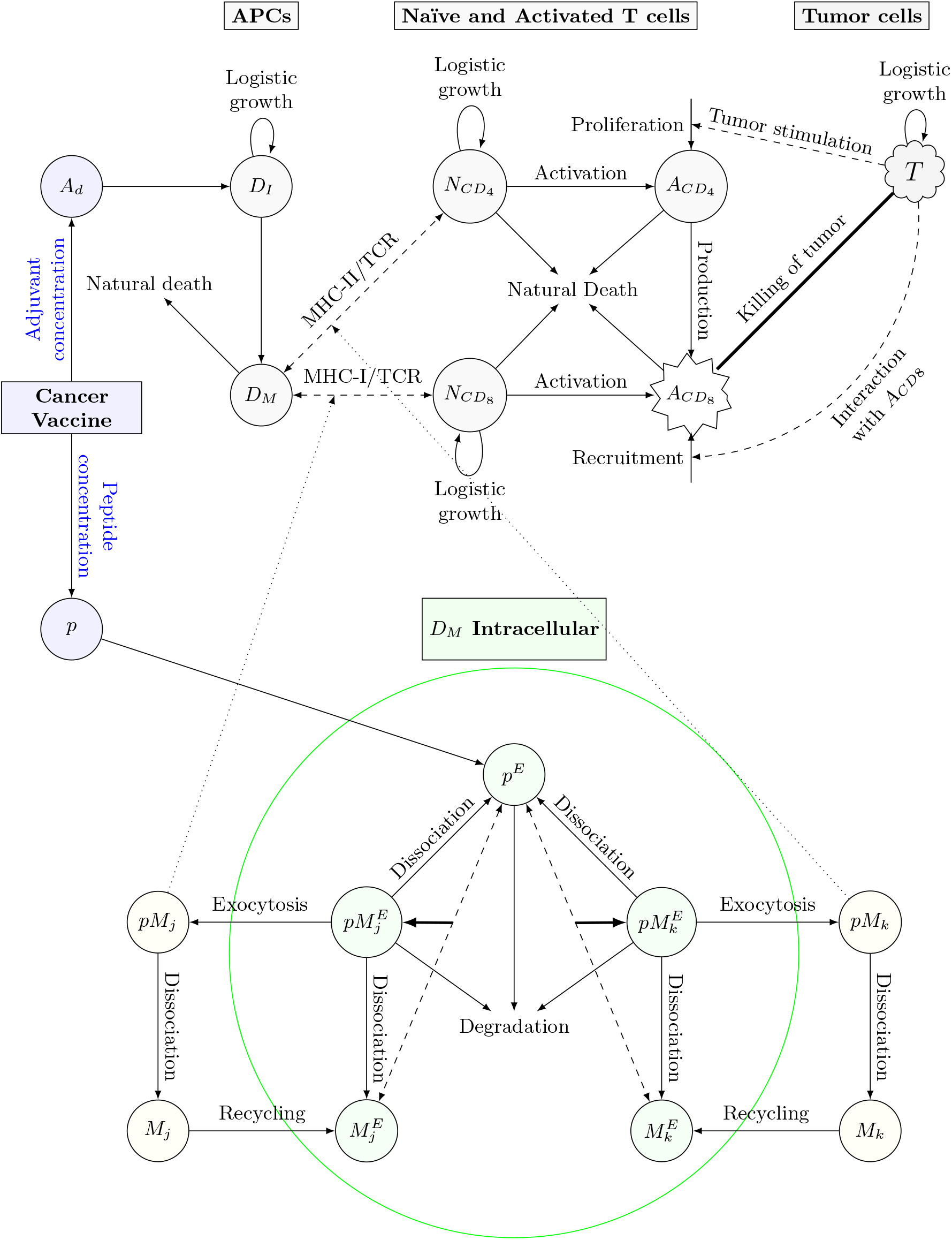
Multiscale flow diagram of immunogenicity to cancer vaccine. The immunological mechanism starts when a patient receives a cancer vaccine, a combination of immunogenic peptides and adjuvant. Adjuvant helps enhance the maturation of immature DCs for antigen presentation at the mature DC surface. Endocytosed peptides interact with MHC-I/II molecules at the subcellular level in matured DCs through binding, dissociating or degradation. Subsequently, antigen-specific T cells are activated by peptides bound to MHC-I/II. Only activated *CD*8^+^ T cells can kill tumor cells. Nonetheless, activated *CD*4^+^ T cells can help activate *CD*8^+^ by tumor stimulation or secretion of IL-2. The solid (dashed) arrows indicate direct (indirect) interactions between the populations at the cellular or subcellular level. The dotted arrows indicated interactions between populations at the subcellular and cellular levels. (APCs: Antigen presenting cells, TCR: T cell receptors. *j* and *k* determine the number of specific MHC-I/II allelic molecules.

**Table 1.** Description of model population (or state) variables including units.

|  | Population | Definition | Units |
| --- | --- | --- | --- |
| **Vaccine** | $p$ | Peptide Concentration | pmol |
| | $A_d$ | Adjuvant Concentration | mg/L |
| **Cellular** | $D_I$ | Immature DCs | cells |
| | $D_M$ | Mature DCs | cells |
| | $N_{CD4}$ | Naïve $CD4^+$ T cells | cells |
| | $N_{CD8}$ | Naïve $CD8^+$ T cells | cells |
| | $A_{CD4}$ | Activated $CD4^+$ T cells | cells |
| | $A_{CD8}$ | Activated $CD8^+$ T cells | cells |
| | $T$ | Tumor cells | cells |
| **Molecular** | $p^E$ | Endosomal peptide fragments | pmol |
| | $M_s^E$ | Free Endosomal MHC-I or MHC-II | pmol |
| | $pM_s^E$ | Endosomal p-MHC-I/II | pmol |
| | $pM_s$ | p-MHC-I/II on mature DC membrane | pmol |
| | $M_s$ | Free MHC-I/II on mature DC membrane | pmol |

\* Subscript $s = j$ or $s = k$ determines the MHC-I or MHC-II molecule, respectively.

In addition to modeling the immunological cascade produced by a personalized cancer vaccine, we used patient-specific data from six patients with melanoma at different disease stages to calibrate the MRM model and estimate key model parameters [27]. Then, we performed *in silico* experiments (virtual simulation) to investigate the response of each patient to cancer immunotherapy and assess changes in tumor sizes.

### Dosing optimization problem

In this study, we formulate two optimization problems associated with the MRM model. We will refer to MRM model as the state system for the purpose of the optimization setup. Our optimization problem is defined based on the settings and patient information from a published clinical trial [27]. The trial consisted of six patients with stage III (Patients 1, 3, 4 and 5) and stage IV (Patients 2 and 6) melanoma who completed a full series of an immunogenic personalized neoantigen cancer vaccine and followed-up for approximately 6 months. Specifically, the treatment consisted of a series of five priming and two booster vaccinations. Each vaccine dose was formulated with a set of immunizing peptides unique for each patient, admixed with an adjuvant (Polyinosinic-polycytidylic acid, and poly-L-lysine (poly-ICLC)).

In order to formulate the dosing-optimization problem for the cancer vaccine model, we use the optimal control theory of ODEs [28]. This theory has been extensively used in the literature to support informative decisions regarding different biological systems [10, 20, 22, 25, 29]. We followed the three main steps to formulate an optimal control problem: (1) define a biological system (e.g., a system of ODEs), (2) define a set of admissible controls, and (3) define an objective functional or target that entails the purpose of the optimization.

Once the optimization problem is defined, we derive a set of necessary conditions that the optimal control as well as the corresponding states must satisfy. Lastly, we use the necessary conditions to numerically solve the optimization problem using the Forward-Backward Sweep Method, as described below.

To formulate our optimal control problem, some assumptions are made. The earlier published MRM model assumed that the vaccine is administrated to a patient as a series of doses as described by the function

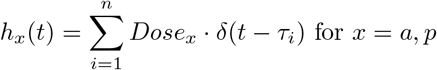

where *τ*_*i*_ represents the day of the vaccine administration with schedule *i* = 1, 2, …, *n*. Moreover, it was also assumed that the values of *Dose*_*x*_ for *x* = *a, p* (*a* for adjuvant and *p* for peptide) were constant, i.e., a patient’s vaccine dose consisted of fixed concentrations of peptide and adjuvant at each administration throughout the treatment. However, in this study, the goal is to find a set of peptide and adjuvant concentrations that is optimal for specified target. We assume the functions *Dose*_*p*_(*t*) and *Dose*_*a*_(*t*) respectively, representing the vaccine concentration composed of peptides (pmol) and adjuvant (mg), are now piecewise functions of time that take nonzero values during the time of vaccination 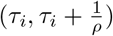, and are zero otherwise. Here, based on the vaccination schedule described in [27], we assume that the vaccine is given to patients at a fixed schedule *τ*_*i*_ = 0, 3, 7, 14, 21, 83, 139 days for *i* = 1, …, 7. Moreover, *ρ* = 0.001 days^*−*1^ or 86.4 seconds, which is an approximation of the time taken for a subcutaneous vaccination process [30].

Based on the above assumptions, our proposed rate of peptide and adjuvant concentrations are described by Equations (1) and (2)

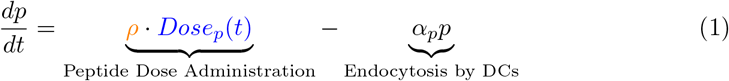

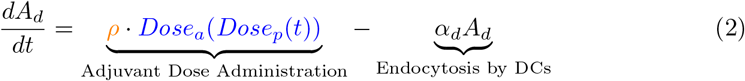

where *α*_*p*_ and *α*_*d*_ are rates of DC uptake for the peptide and the adjuvant molecules, respectively. Reflecting the synergy between these two vaccine components, the amount of adjuvant is determined by a fixed adjuvant:peptide ratio and the amount of peptide in mg

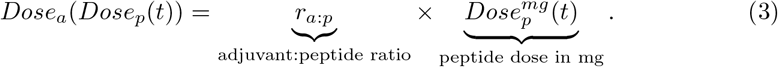

Where

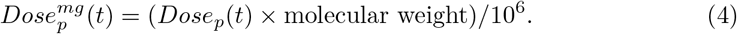

converts the pmol concentration of peptides to units in mg. With the assumption that the adjuvant amount depends on the amount of peptides, we will always guarantee that both vaccine components are present in each vaccine dose. The adjuvant, as an immunostimulatory agent, activates the DCs and lead to their maturation. Peptides are trafficked to the endoplasmic reticulum and endosome of mature DCs, interacting with MHC class I and II molecules.

In this particular case, we assume a state system (MRM model) with the above assumptions and a set of acceptable peptide concentrations per unit volume:

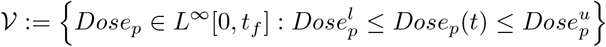

where *t*_*f*_ is the length of time for the treatment. The lower and upper bounds, 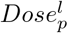 and 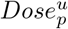, refer to the allowed minimum and maximum concentrations of peptide. It is assumed that the lower bound, 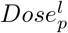, is positive because negative values are not practically meaningful. The total number of vaccination doses administered to a patient throughout the whole therapy is *τ* ; where *τ* = 7 in our case study. The *L*^*∞*^[0, *t*_*f*_] notation is the space of all bounded functions in the interval [0, *t*_*f*_].

We propose the following two objective functionals:

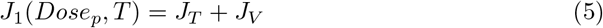

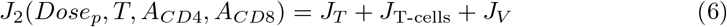

where

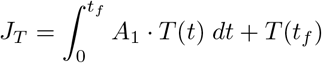

includes two terms, the residual tumor cells over the course of the treatment, 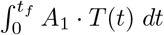, and residual tumor cells at the final time, i.e., *T* (*t*_*f*_). The integral

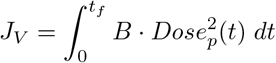

measures the total amount of peptide concentration and

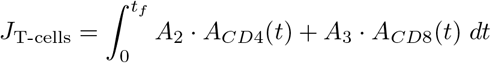

represents the total number of activated T cells (*A*_*CD*4_ and *A*_*CD*8_) from the beginning to the end of the therapy. Note that by minimizing the peptide concentration through *J*_*V*_, the vaccine dose (including peptide and adjuvant) is implicitly minimized since we set a fixed ratio for adjuvant:peptide.

Both objective functionals *J*_1_ and *J*_2_ share the terms *J*_*T*_ and *J*_*V*_, but *J*_2_ has an additional term, *J*_T-cell_. This means that both objective functionals target high tumor killing and low vaccine concentrations, but *J*_2_ integral additionally targets to minimize the excess T cell response, which may adversely affect the safety of the treatment. An excessive T cell response could cause autoimmune reactions and tissue damages in patients [31, 32]. Note that the term *J*_*V*_ has value of 0 except for a short period of time (assume to be the average time of injection, *ρ*^*−*1^). The constants *A*_1_, *A*_2_, *A*_3_ and *B* are weight parameters that measure the relative importance of each integral to the number of tumor cells at the end of therapy, *T* (*t*_*f*_). These parameters were manually calibrated for each patient by model simulations using their longitudinal T cell data. The units of these weight parameters are shown in Table 2.

**Table 2.** Weight parameter units

| Weight Parameter | Units |
| --- | --- |
| $A_1, A_2$ and $A_3$ | 1/time |
| $B$ | cells/pmol<sup>2</sup> × 1/time |

Moreover, we establish the total immune and tumor responses by calculating the areas under the curve (AUC) of the time series of activated T cell populations (*A*_*CD*4_ + *A*_*CD*8_), and the tumor cell population (*T*), respectively.

### Optimal dosing problem formulation

We illustrate the optimal dosing problem setup using the objective functionals *J*_1_ and *J*_2_. The optimal dosing problem consists of

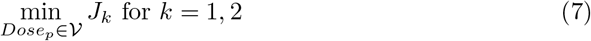

subject to the cancer vaccine immunotherapy model, i.e., the state system with aforementioned assumptions, and non-negative initial conditions. Therefore, our optimization problem is a minimization problem.

The goal of the minimization problem when using *J*_1_ is to find a peptide minimum concentration, 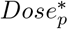 (not necessarily unique), along the entire duration of the cancer immunotherapy so that the residual tumor cells are minimized. For *J*_2_, the goal is to minimize activated T cell count in addition to the residual tumor cells. We derive the necessary conditions of the minimization problem as described in S1 Appendix.

To determine the efficacy of a vaccine dose, we use the objective functionals to compare the “score” of an optimal vaccine dose against any other acceptable vaccine dose (e.g., the vaccine dose given to patients during the clinical trial and other selected multipliers). When the following inequality holds at an optimal vaccine dose

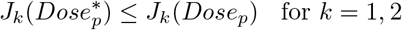

the ratio

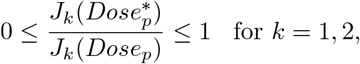

determines the efficacy of an optimal vaccine dose against another acceptable vaccine dose. When this ratio is close to 1, the other acceptable vaccine dose provides similar outcomes as the optimal vaccine dose, while when this ratio is close to 0, the other acceptable vaccine dose is far from achieving the desired goal, i.e., to minimize the total tumor size or total tumor size and immune response, as compared to the optimal vaccine dose determined by our model and simulations.

However, since there is no guarantee that 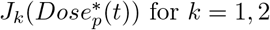 for *k* = 1, 2 is a global minimum, we can also have that for some *Dose*_*p*_ ∈ 𝒱

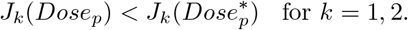

In this case, the ratio

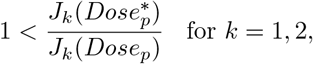

determines the inefficacy of an optimal vaccine dose to minimize an objective functional. When this occurs, we say the performance of the optimal vaccine dose is worse than the other acceptable vaccine dose. Thus, it will be better to choose the other acceptable vaccine dose, *Dose*_*p*_(*t*), or try to refine the set of weight parameters to obtain a different optimal vaccine, 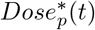.

In the next section, we discuss the numerical approach to solve the minimization problem.

### Forward Backward Sweep Method (FBSM)

The FBSM is an iterative algorithm to solve optimal control problems. In general terms, the numerical scheme consists of solving two sets of coupled differential equations and using the optimal control characterization to update the new solutions. The method exits the loop once the desired convergence criteria for solutions are achieved. The FBSM has been used extensively to solve optimal control problems involving biological, immunological, and ecological systems (ODEs, Partial Differential Equations, Difference Equations, Delayed Equations, Integro-differential Equations, etc.) [28]. For details on numerical convergence and stability of the method, see [33].

In particular, we implement the FBSM to numerically find the solution to the dosing-optimization problem. The state system is solved forward in time while the adjoint system is solved backward in time using initial conditions for the state variables and *transversality condition* for the adjoints. Different ODE solvers, such as `ode45` in Matlab or `solve_ivp` from `scipy.integrate` library in Python, can be used to implement this routine. Additionally, we use the optimal dosing characterization to update and find an optimal set of peptide and adjuvant concentrations. The state and adjoint systems with the optimal dosing characterization can be found in S1 Appendix.

The codes to reproduce the results shown in this paper are available in the following GitHub repository https://github.com/Wenvalegam/CanVaxDOpt_Model. A flow chart summarizing key steps to apply the FBSM to solve our optimization problem is shown in Fig 2.

**Fig 2.**
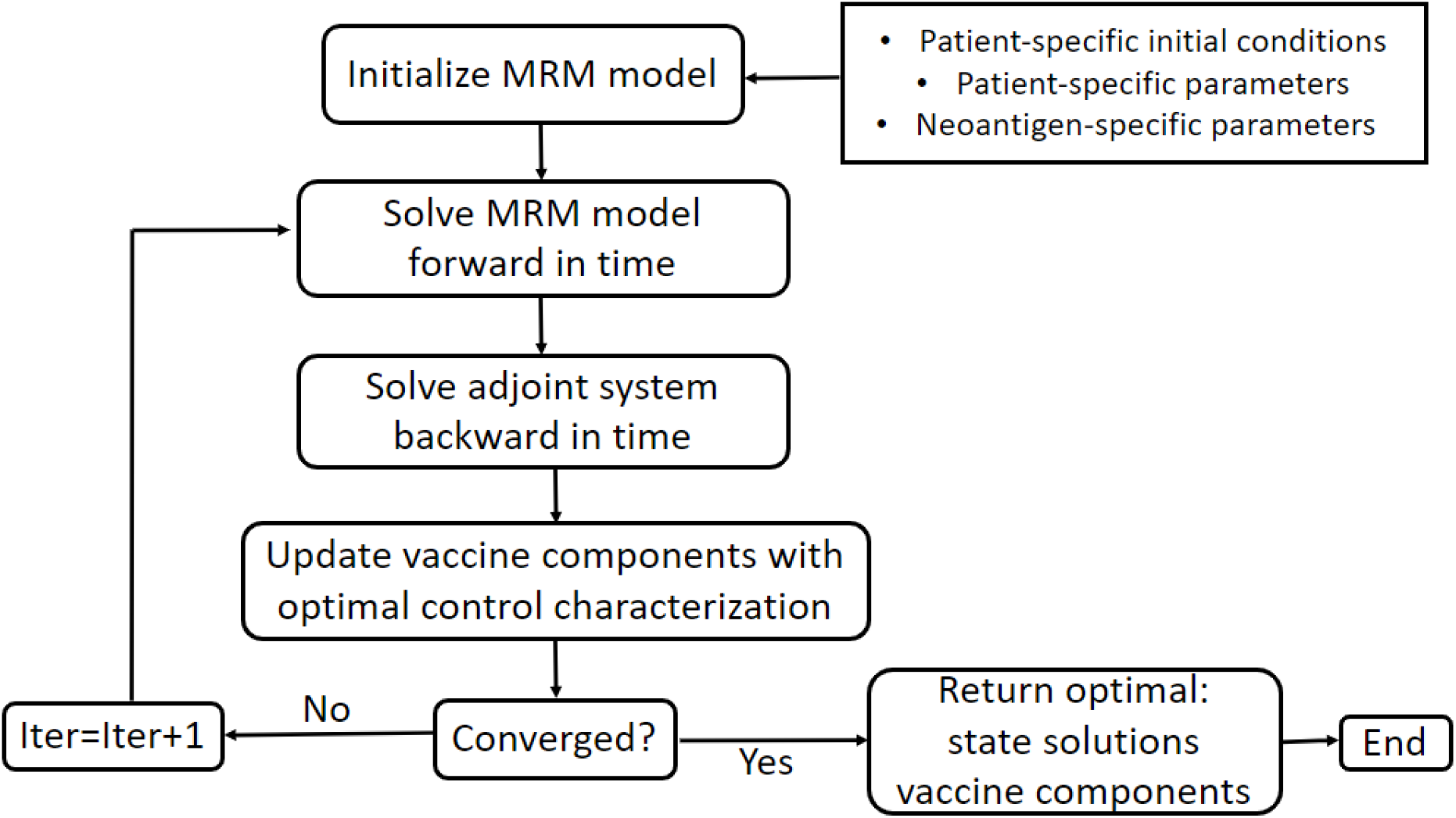
Flow chart of the FBSM. Patient-specific initial conditions are the values of all population variables in Table 1 at the start of the treatment. Patient-specific parameters are T cell recruitment and tumor killing rates. Neoantigen-specific parameters refer to unique immunogenic peptide sequences and binding affinities. All these values can be found in MRM paper for each patient in our case study.

## Results

### Dosing optimization for six patients with melanoma

In practice, it is very difficult to optimize the dose of a cancer vaccine for individual patients. A dose is typically determined and given to all patients, but some questions remain. For instance: Is the dose provided to an individual patient optimal? Is a fixed dose for all vaccinations warranted or should the doses for vaccinations over the treatment vary? The dosing optimization problem is designed to understand the effects produced by a personalized neoantigen cancer vaccine in six patients with melanoma when the concentrations of peptide and adjuvant are varied. First, we formulate a minimization problem to quantify the impact of the components of the cancer vaccine, namely, peptide and adjuvant (in a fixed ratio), on the total number of tumor cells (*T*) of each patient. With this optimization exercise, we can understand how effective the vaccine is on each patient by predicting tumor size reductions through model simulation. A second optimization problem evaluates the effect of vaccine dose on the total number of tumor cells and an addition of activated T cells (*A*_*CD*4_ + *A*_*CD*8_). In this case, we optimize the vaccine dose based on achieving the minimum number of tumor cells (effectiveness) and activated T cells (safety), identifying a vaccine dose that elicits an immune response strong enough to minimize the tumor size, but not in excessive amount to cause safety concerns. Such vaccine dose would have the optimal benefit-risk profile.

To solve the optimization problem, we initialize the MRM model (or state system) using the set of parameters described previously [26] (Table 1 and Table S1). The concentrations of neoantigen peptides used in the clinical trial for each patient are presented in Table 3. For more details on how these values are derived, see the supplemental information in [26]. In the clinical trial, each peptide pool (four pools per vaccine) was admixed with 0.5 mg of Poly-ICLC adjuvant in a volume of 1 ml of aqueous solution. The adjuvant is added to the vaccine formulation to enhance immunogenicity [34]. For instance, on each vaccination day, Patient 1, received a cancer vaccine dose including 3.9 mg of peptides and 2 mg of adjuvant (poly-ICLC) in a volume of 4 ml aqueous solution [27]. The adjuvant:peptide ratio, *r*_*a*:*p*_, used in clinical trial for each patient can be found in Table 3, which are also used as fixed ratio in our model simulations for each patient.

**Table 3.** Peptide dose and adjuvant:peptide ratio converted from clinical trial data for patients with melanoma [26].

| Patient | No. of peptides | Weight (mg) | Amount (pmol) | Adj:Pep ($r_{a:p}$) |
| --- | --- | --- | --- | --- |
| 1 | 13 | 3.9 | 119,340 | $\frac{2}{3.9}$ |
| 2 | 17 | 5.1 | 120,030 | $\frac{2}{5.1}$ |
| 3 | 14 | 4.2 | 109,570 | $\frac{2}{4.2}$ |
| 4 | 14 | 4.2 | 116,570 | $\frac{2}{4.2}$ |
| 5 | 20 | 6 | 110,860 | $\frac{2}{6}$ |
| 6 | 20 | 6 | 111,820 | $\frac{2}{6}$ |

We assumed that the average duration of the clinical trial including post-vaccination follow-up is 200 days. The set of acceptable peptide concentrations per 4 ml of aqueous solution is determined by

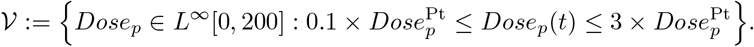

The 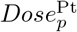 is the peptide dose used in the clinical trial for each patient and is presented in the 4^th^ column of Table 3. We vary the peptide concentrations from 0.1-fold to up to 3-fold of the 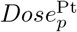 in our model simulations. Note that there is no need to select the same lower and upper fold-bounds of peptide concentrations for all the patients, but here we adopt this approach for numerical simplicity.

The exact value for the adjuvant dose can be found using Equation (3) with the last column in Table 3 and the corresponding *Dose*_*p*_(*t*) in mg. We apply the dosing-optimization problem to each of the six patients within their set of acceptable doses (𝒱) to predict outcomes of using an optimal vaccine dose and other suboptimal doses including the dose used in the clinical trial and their 0.1-3 folds deviations.

The solutions of the vaccine dosing minimization problem provide us with an optimal concentration of peptide denoted by 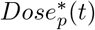 to achieve either *J*_1_ or *J*_2_ objectives. In particular, to determine an optimal total vaccine dose that a patient should receive on a vaccination day, we compute the following integral

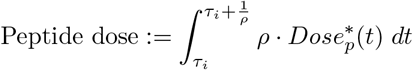

where *ρ*^*−*1^ is the average time of injection and *τ*_*i*_ for *i* = 1, 2, …, 7 corresponding to a vaccination day, and thus, the number of doses a patient will receive. The optimal total dose of peptide administered over the whole treatment is computed with the following integral

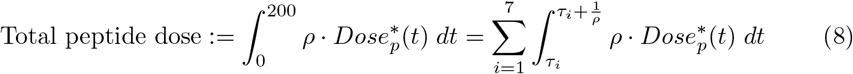

which constitutes the contribution of peptide from each vaccination day. As noted earlier, we assume *Dose*_*p*_(*t*) is 0 outside the vaccine administration periods, 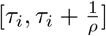 for *i* = 1, 2, …, 7, to derive the formula in Equation (8).

The model’s non-specific patient parameters are taken from Table S1 in [26]. Note that the maximum activated T cell recruitment rates (for *A*_*CD*4_ and *A*_*CD*8_), *c*_4_ and *c*; maximum lysis rate by activated T cells, *d*; and the dependence of lysis rate between T cells and tumor, *λ*; are all patient-specific parameters. The values for these parameters are taken from Table 1 in [26], where authors used a global optimization tool to find parameters best fit to individual patient’s data (with adjusted *R*^2^ between 0.75 and 0.95). Moreover, the off rate of peptide-MHC type I/II with allele *s, k*_off,*s*_ for *s* = *j, k*, are neoantigen specific parameters, which can be accessed using GitHub repository https://github.com/Wenvalegam/CanVaxDOpt_Model.

### Immune response

The model predicts the number of activated T cells (*A*_*CD*4_ + *A*_*CD*8_) over a 200 day period for six patients applying an optimal vaccine dose, clinical trial dose, and half of the clinical trial dose. The results are plotted in Fig 3 (for *J*_1_) and 4 (for *J*_2_).

**Fig 3.**
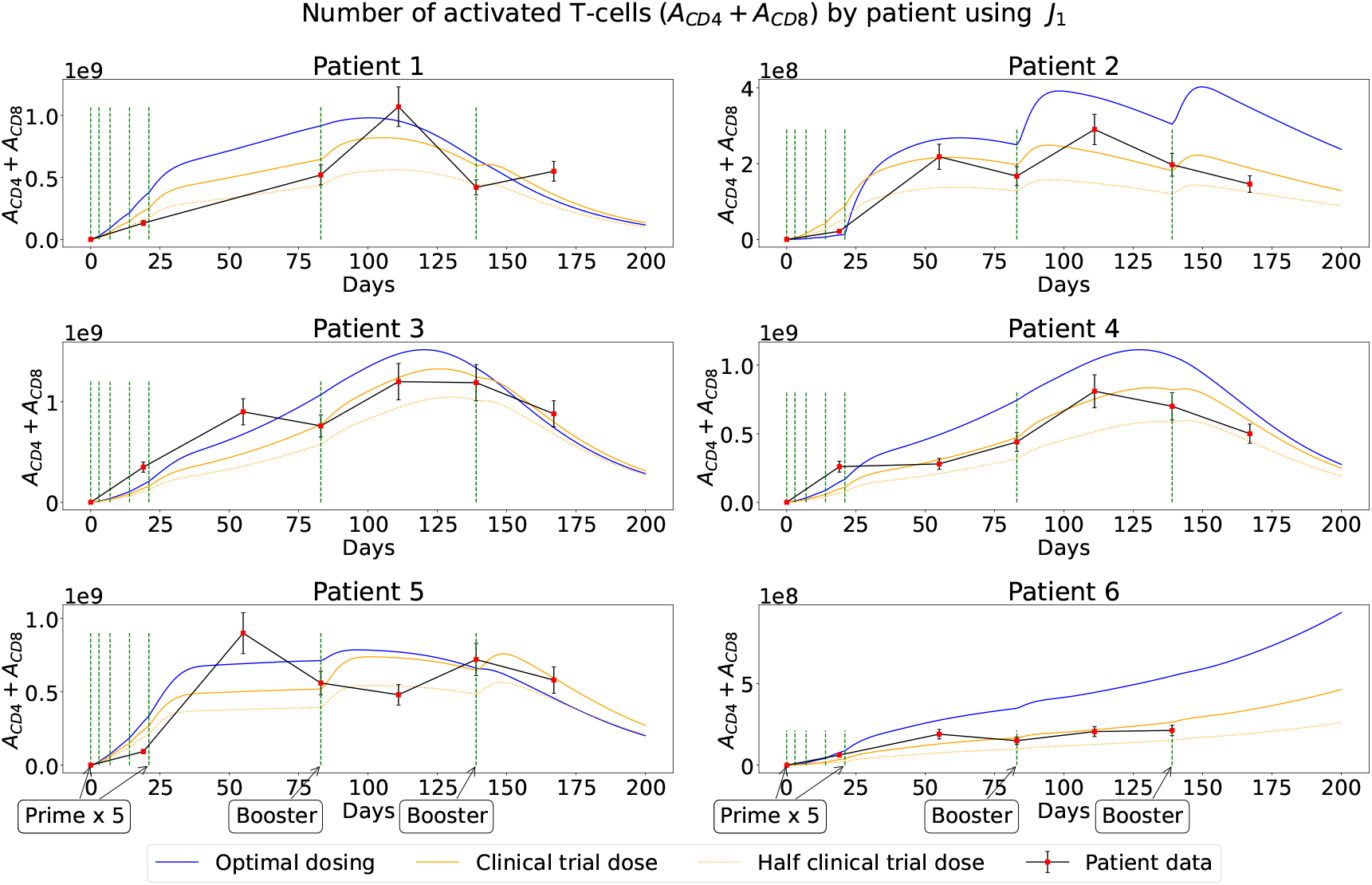
Activated T-cells (*A*_*CD*4_ + *A*_*CD*8_) when optimal (blue) and suboptimal (dark orange for clinical trial dose and dotted orange for half clinical trial dose) are applied to each patient using *J*_1_. Red dots represent patients’ measurements at specific times in clinical trial with 15% standard error. The vertical green dashed lines correspond to the days of vaccination.

In Fig 3, we can see that, overall, the optimal vaccine dose (from *J*_1_) produced a stronger immune response (higher cell count of *A*_*CD*4_ + *A*_*CD*8_) than any other tested vaccine dose for all patients. On the other hand, in Fig 4, we observe that the immune response of almost all the patients with optimal vaccine dose (from *J*_2_) was significantly lower (lower cell count of *A*_*CD*4_ + *A*_*CD*8_) than their immune response to any other dose. These observations demonstrate the importance of establishing objective functionals, which are used to select an optimal vaccine dose for an individual patient.

**Fig 4.**
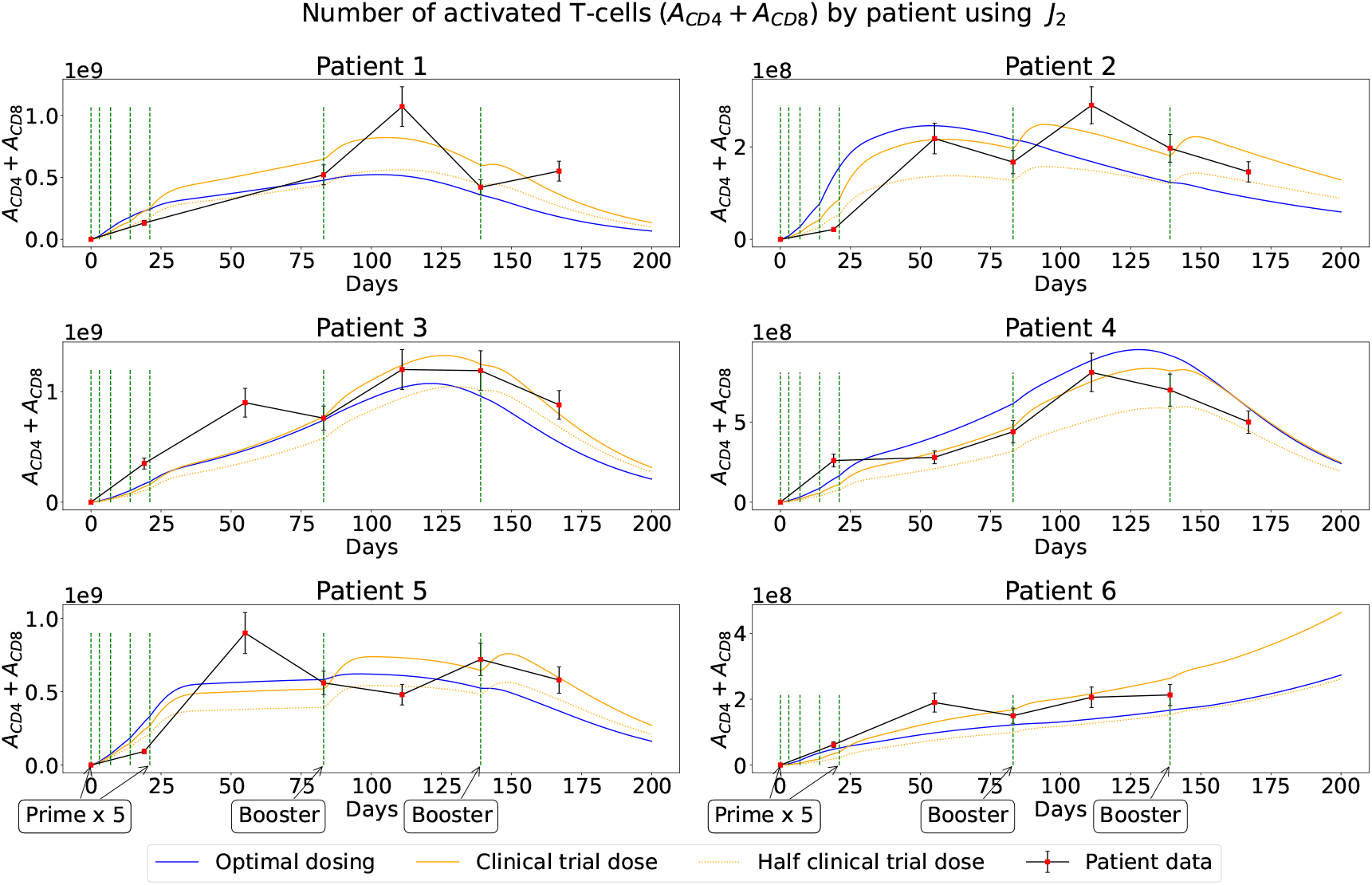
Activated T-cells (*A*_*CD*4_ + *A*_*CD*8_) when optimal (blue) and suboptimal (dark orange for clinical trial dose and dotted orange for half clinical trial dose) are applied to each patient using *J*_2_. Red dots represent patients’ measurements at specific times in clinical trial with 15% standard error. The vertical green dashed lines correspond to the days of vaccination.

### Tumor response

In Fig 5, we depict the evolution of the tumor size in mm over 200 days for each patient under optimal and suboptimal vaccine dosing. Note that we first obtained the number of tumor cells and then converted the cell number into mm by using the diameter formula derived in [26]

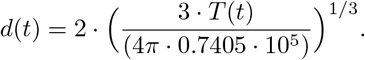

Moreover, the tumor responses for most of the patients based on *J*_2_ (not shown) are similar to the cases for *J*_1_ shown in Fig 5, with the exception of Patient 2 and 6. When optimizing *J*_2_, these two patients have a larger tumor size on the day 200 (16.38 mm and 6.4 mm, respectively). In general, we observe that the optimal vaccine dose (blue line) performed slightly better than suboptimal vaccines within the vaccination period (day 0 and 139) of the immunotherapy for all patients, but Patient 2. However, at the late stage, all tested vaccine doses show similar effectiveness in reducing the tumor size for all the patients with the optimal vaccine dose performing noticeably better for Patient 6.

**Fig 5.**
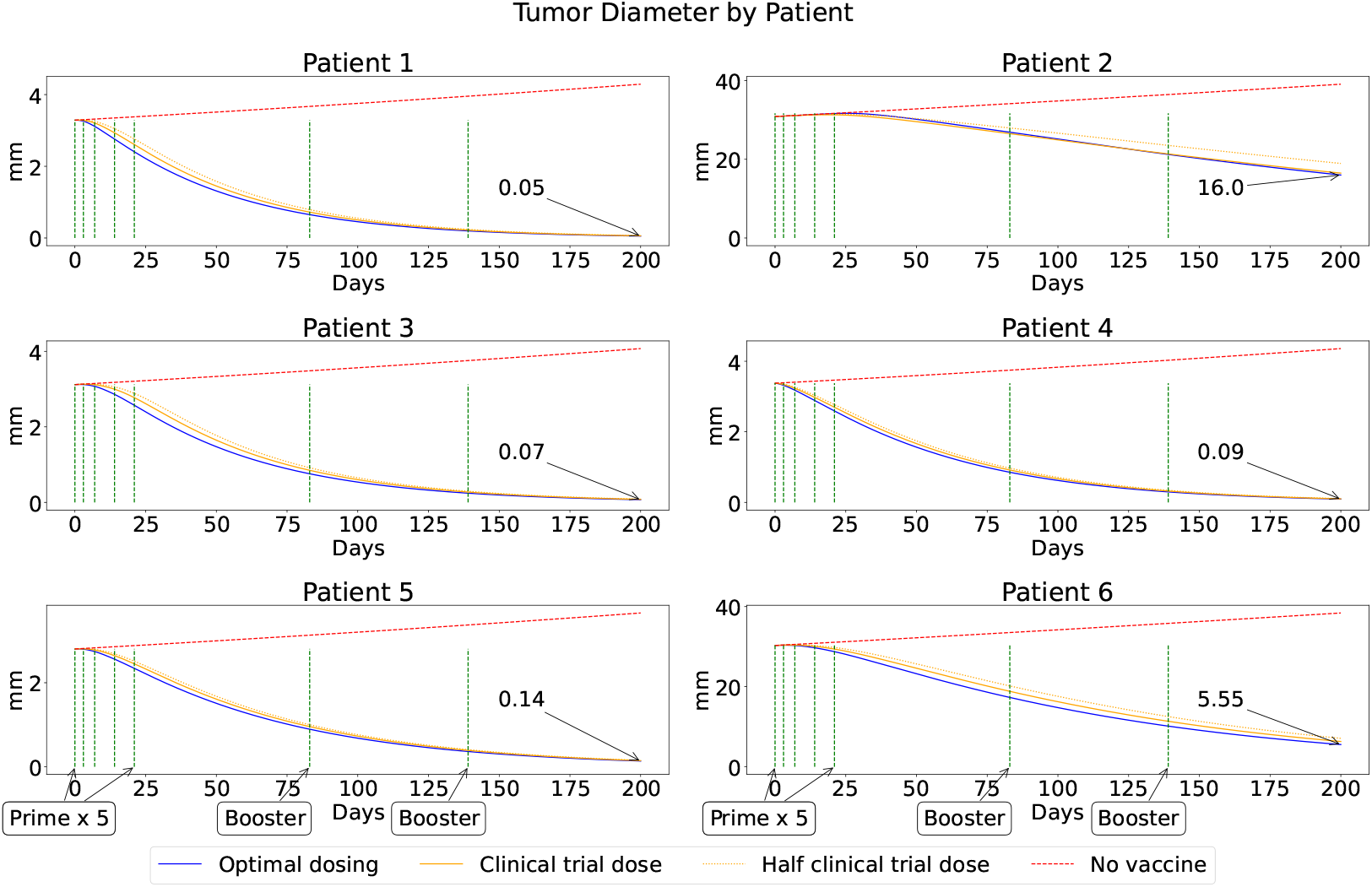
Tumor diameter when optimizing *J*_1_ and optimal (blue), suboptimal (clinical trial dose and half of clinical trial dose represented by dark and light orange, respectively) and no (red) vaccinations are applied. Numbers with an arrow pointing at the blue curve when *t* = 200 days, represent the number of tumor cells at the end of the treatment. The vertical green dashed lines correspond to the days of vaccination.

### Vaccine doses

The optimal dose concentrations of peptide and adjuvant for each vaccination of six patients identified through our simulations are shown in Fig 6 (for *J*_1_) and 7 (for *J*_2_). These concentrations were used to obtain the optimal tumor responses (blue curves) in Fig 3 (*J*_1_) and 5 (*J*_1_), and the optimal immune and tumor responses in Fig 4 (*J*_2_). The exact values for these doses can be found in S1 Appendix. The optimal concentrations of peptides and adjuvants when considering minimizing the tumor response only (*J*_1_) are demonstrated by Fig 6. In this case, for patients 1, 3, 4, 5 and 6 the levels of peptide and adjuvant as 3-fold of clinical trial dose (the upper bound of prespecified range) for primes are optimal. For Patient 2, approximately 0.1-fold of the clinical trial dose at the first four vaccinations, while 3-fold of clinical trial dose for the subsequent vaccinations are optimal. This particular situation of Patient 2 implies that starting time point of cancer vaccine may be different among the patients. With the optimal dose obtained for *J*_1_, all patients achieve tumor size reduction similar to that reported in clinical trial [27] and are consistent with model predictions [26]. The optimal concentrations of peptides and adjuvants when considering minimizing the T cell and tumor responses (*J*_2_) for each patient are shown in Fig 7. Interestingly, our results show that all patients need higher doses for the initial several primes, but lower doses afterward. This observation suggests that when trying to minimize the immune response in addition to the tumor response (*J*_2_) the initial vaccination dose should be higher; while lower afterward.

**Fig 6.**
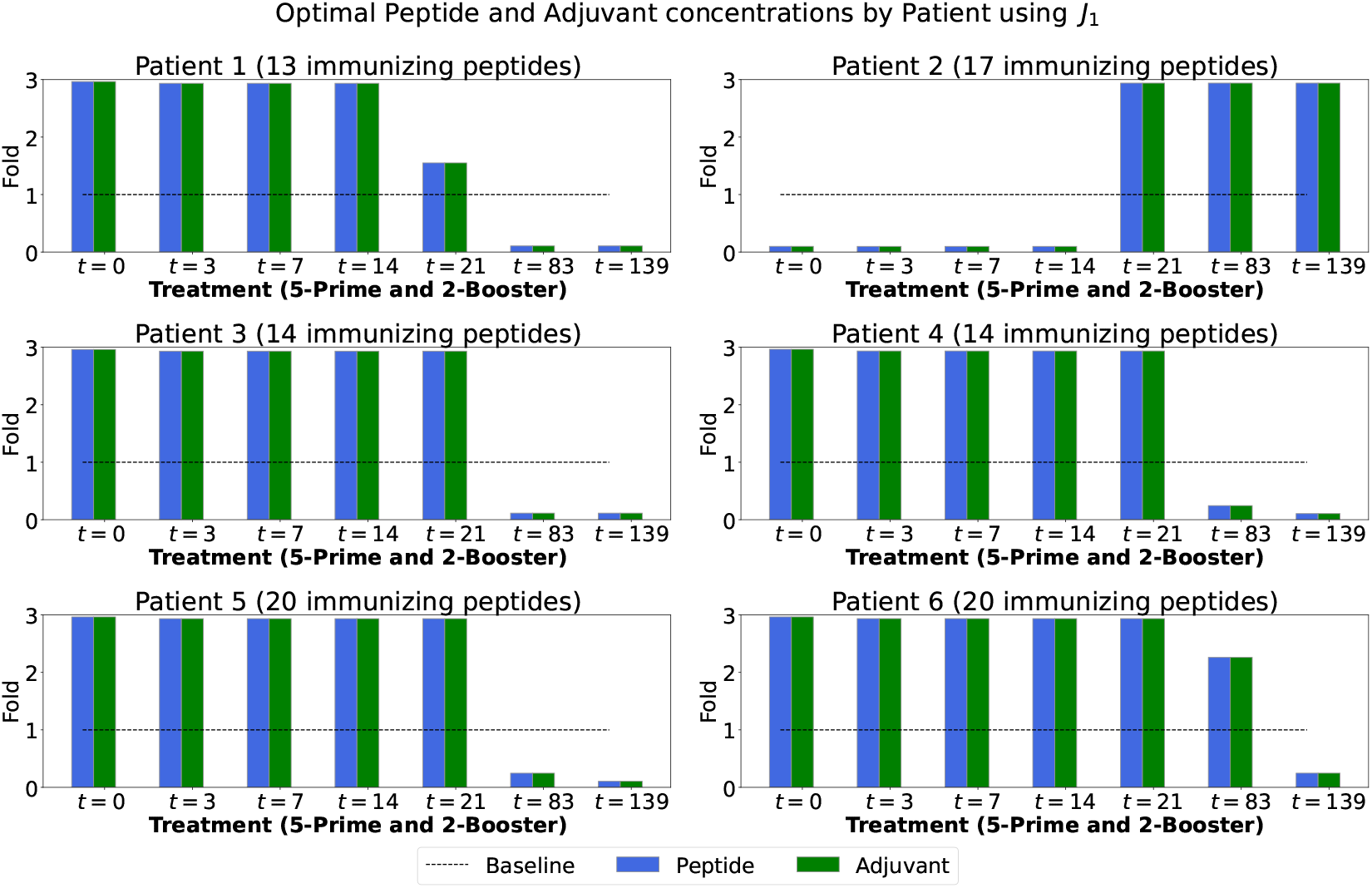
Bar plots correspond to optimal peptide and adjuvant (poly-ICLC) doses as the number of folds of clinical trial dose for each vaccination using *J*_1_. The horizontal dashed line represents the dose used in the clinical trial (baseline) given in Table 3 for each patient. The total number of immunizing peptides is also reported at the top of each panel.

**Fig 7.**
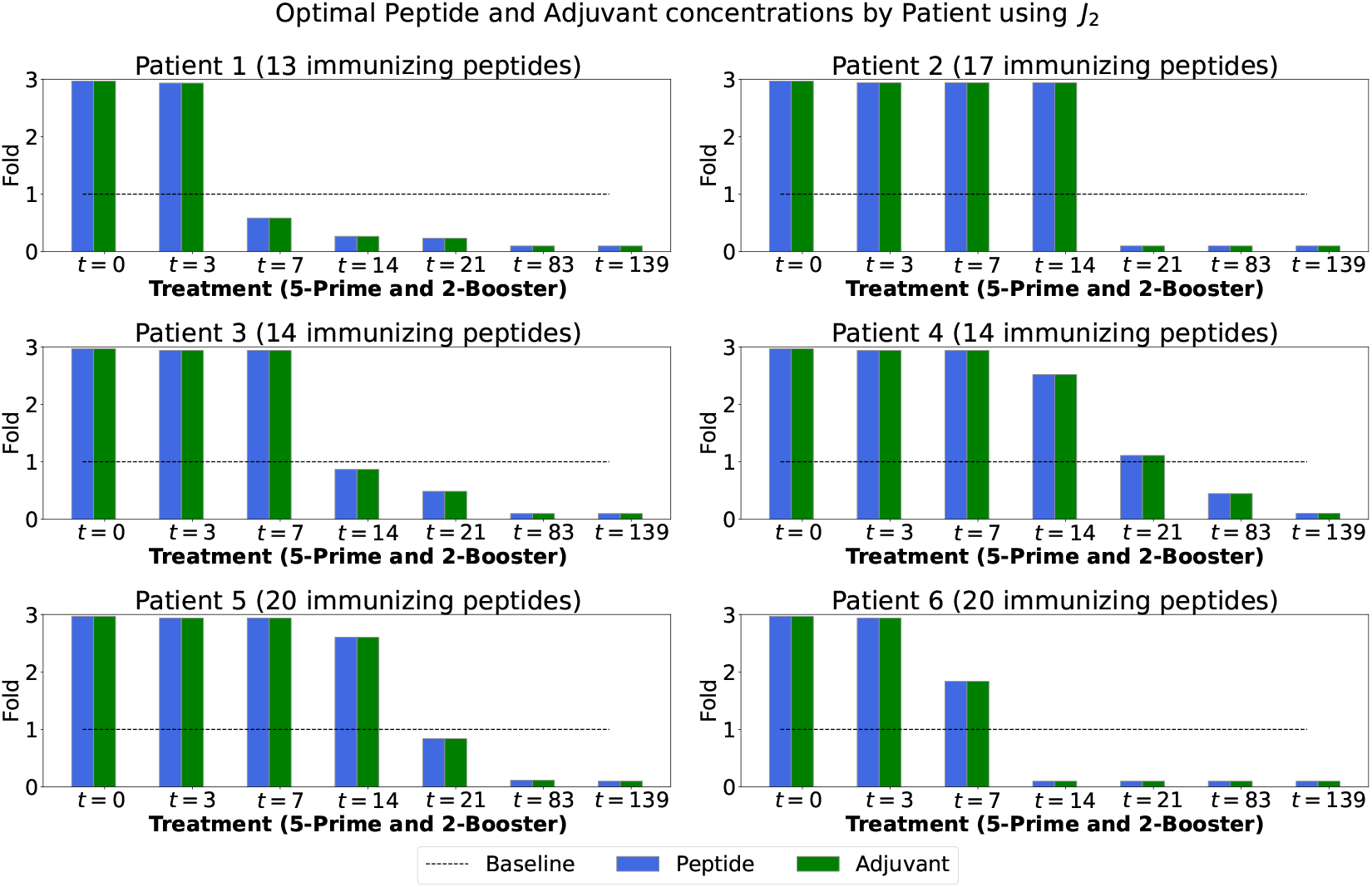
Bar plots correspond to optimal peptide and adjuvant (poly-ICLC) doses as the number of folds of clinical trial dose for each vaccination using *J*_2_. The horizontal dashed line represents the dose used in the clinical trial (baseline) given in Table 3 for each patient. The total number of immunizing peptides is also reported at the top of each panel.

### Immune and tumor responses to vaccine doses

The total immune (sum of activated T cells) and total tumor responses (sum of tumor cells) to optimal and suboptimal vaccine doses over the total length of the immunotherapy are shown in Fig 8 and 9 when objective functionals *J*_1_ and *J*_2_ are minimized.

**Fig 8.**
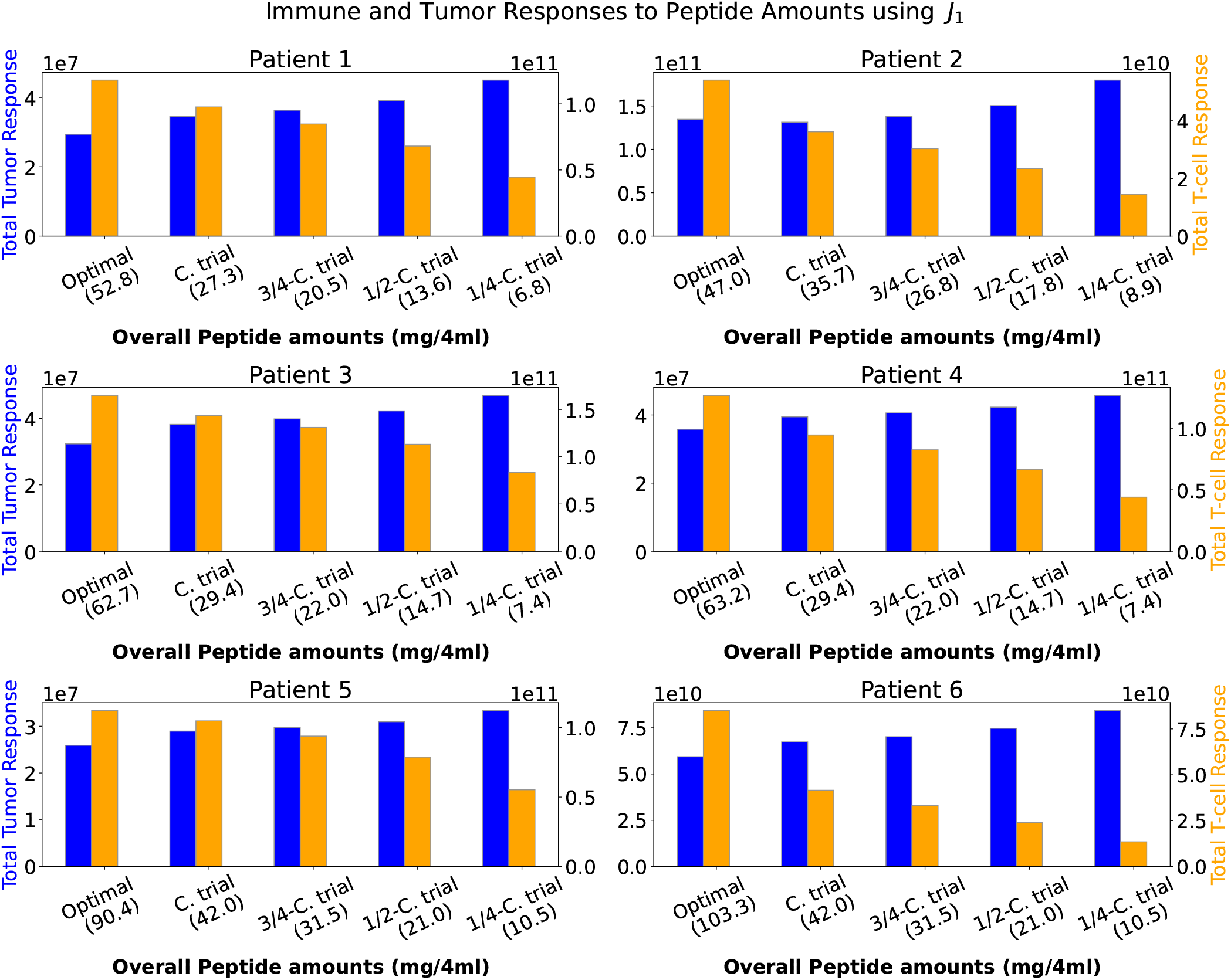
Total immune and tumor responses to optimal and suboptimal vaccine doses using objective functional *J*_1_. Blue bars represent the sum of tumor cells over treatment period with vertical axis to the left. Orange bars correspond to sum of activated T cells over treatment period with vertical axis to the right. C. trial stands for clinical trial dose. The peptide doses along the horizontal axis correspond to the total peptide amounts that each patient received over treatment in relation with clinical trial dose when applying optimal or suboptimal vaccine doses. Blue and orange bars were calculated by computing the AUC over [0, 200] of the functions *T* and *A*_*CD*4_ + *A*_*CD*8_.

**Fig 9.**
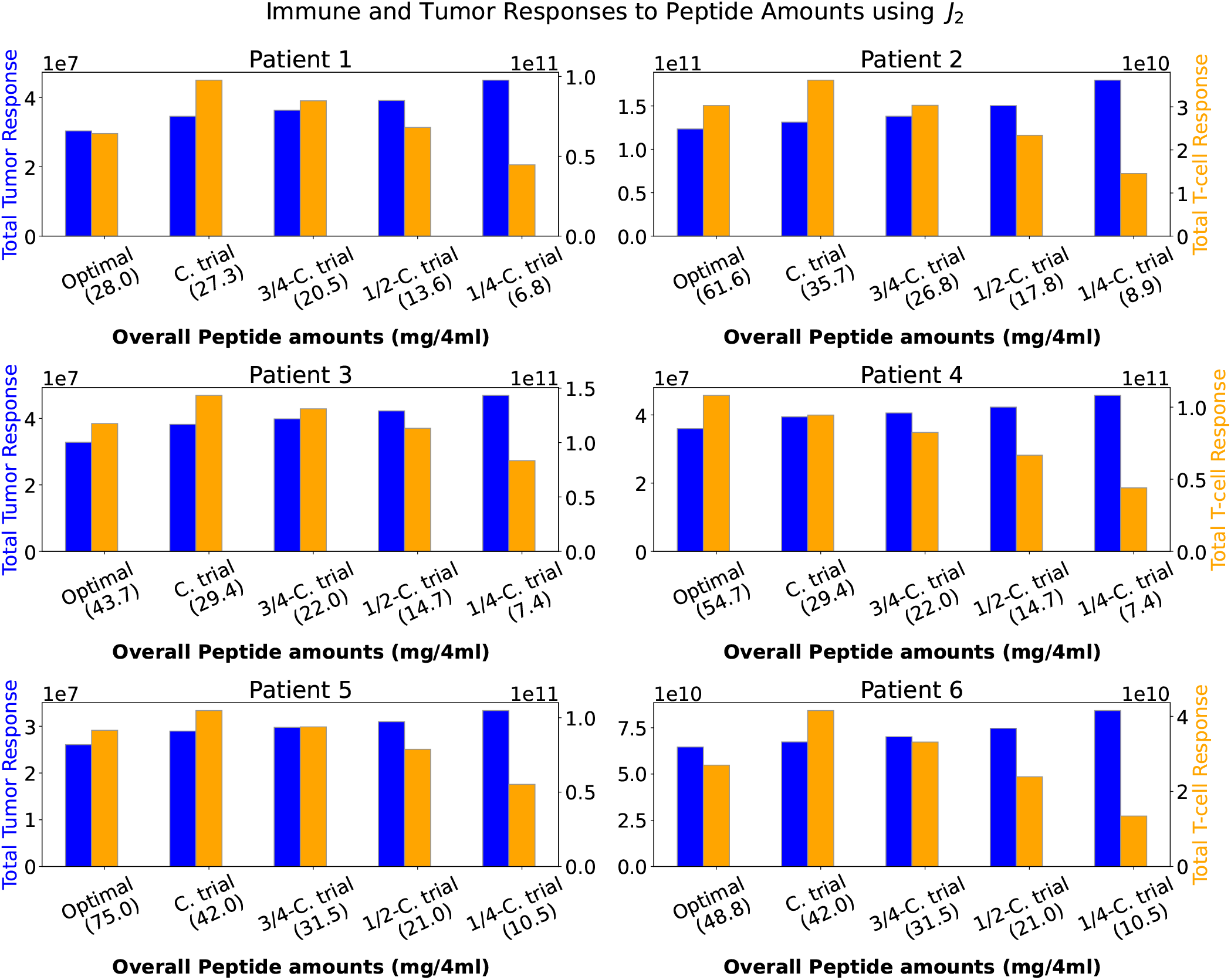
Total immune and tumor responses to optimal and suboptimal vaccine doses using objective functional *J*_2_. Blue bars represent the sum of tumor cells over treatment period with vertical axis to the left. Orange bars correspond to sum of activated T cells over treatment period with vertical axis to the right. C. trial stands for clinical trial dose. The peptide doses along the horizontal axis correspond to the total peptide amounts that each patient received over treatment in relation with clinical trial dose when applying optimal or suboptimal vaccine doses. Blue and orange bars were calculated by computing the AUC over [0, 200] of the functions *T* and *A*_*CD*4_ + *A*_*CD*8_.

The total vaccine doses display at the horizontal axis of Fig 8 and 9 are computed by summing up the terms in Equation (8). For example, the total clinical trial peptide dose amount that Patient 1 received was

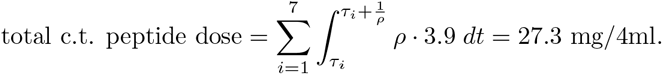

By summing up the total peptide amounts above, the total clinical trial peptide dose for Patient 1 was 41.3 mg/4ml. Similarly, the total optimal or any other tested peptide doses can be computed for other patients. We converted the pmol concentration to mg and plotted them in the figures. Along the left and right vertical axes, we plot the cumulative tumor and immune responses, respectively, by computing the AUC of *T* (*t*) and *A*_*CD*4_(*t*) + *A*_*CD*8_(*t*) along the duration of the immunotherapy associated with each vaccine dose.

In Fig 9, the immune and tumor responses were both minimized (*J*_2_). As shown in Fig 8 for *J*_1_, we see that for most patients the optimal vaccine dose produced a stronger immune response (orange bar) compared with other suboptimal doses. In contrast, in Fig 9 (for *J*_2_), a weaker immune response (orange bar) was observed among all patients, but Patient 4, at the optimal vaccine dose. In both figures, the optimal vaccine dose achieves the same or lower cumulative number of tumor cells (blue bar) when compared to other suboptimal vaccine doses. We notice with either *J*_1_ and *J*_2_ the optimal vaccine doses are effective at overall tumor control or reduction in Patients 1, 3, 4 and 5 over the treatment period. However, for Patients 2 and 6, who have more advanced cancer, significantly greater tumor reduction was achieved when minimizing *J*_1_ than *J*_2_. This result might be true in clinical practice since it suggests that for cancer patients with stage IV melanoma, we may need to focus more on tumor reduction in order to stop, slow and shrink tumor metastasis.

## Conclusion and discussion

Studying the different doses of neoantigen cancer vaccine components that are effective in reducing the residual tumor cells in individual patients, is the aim of this paper. We formulated two mathematical dose optimization problems using optimal control theory. The dose optimization problems were developed to find the optimal vaccine compositions, amount of peptides and adjuvant, that a patient’s vaccine requires to minimize two objective functionals given fixed peptide:adjuvant ratio and vaccination schedule. The MRM model describing the immunological responses at the cellular and subcellular levels are used as the state system for these optimization problems. The first objective functional, *J*_1_, is intended to minimize the amount of peptide dose, and the total number of tumor cells. The second objective functional, *J*_2_, incorporates an additional term of T cells (representing the immune response to the vaccine) in the minimization problem. The necessary conditions that the optimal control problem must satisfy, such as the adjoint system and optimal control characterization, were derived in S1 Appendix.

The vaccine dose optimization problem was applied to the set of six patients from an advanced melanoma cancer clinical trial [27]. We highlighted the immune and tumor responses of each patient to the vaccine doses when minimizing either *J*_1_ or *J*_2_ (Fig 3, 4 and 5). Moreover, we showed the optimal vaccine dose amounts associated with each minimization problem and for each patient, according to the fixed vaccination schedule (Fig 6 and 7). We also illustrated the cumulative immune and tumor responses to the optimal and suboptimal vaccinations over the entire course of the immunotherapy for each patient (Fig 8 and 9).

Although the immune response to the optimal vaccine dose of each patient showed to be reduced significantly when the objective functional *J*_2_ was used instead of *J*_1_, we observed that all patients with stage III melanoma (Patients 1, 3, 4 and 5) have a similar tumor response regardless of which objective functional was used. Patients with stage IV melanoma (Patients 2 and 6) benefited more from using an optimal vaccine dose, when only the tumor response (*J*_1_) was minimized, since greater tumor reductions were observed in comparison to any other tested vaccine dose. The optimal vaccine dose that each patient received minimized the objective functionals *J*_1_ or *J*_2_ when compared to any other tested vaccine doses (as shown in Tables 1 and 2 in S1 Appendix. The additional minimization of the total number of activated T cells in *J*_2_ illustrated that the optimally estimated vaccine doses produced similar effects in reducing the total tumor size.

We observe from *in silico* results that with the help of an optimal vaccine, the tumors of Patients 1, 3, 4 and 5 could possibly be eliminated. For Patients 2 and 6, cancer immunotherapy may reduce the size of the tumor; however the treatment may not be sufficient to move these patients to a less advanced cancer stage as the model predicted a large tumor size at the end of the treatment period. Other combination treatment options, such as radiotherapy or immune checkpoint inhibitors, may improve these patients’ quality of life by overcoming tumor persistence and increasing clinical efficacy [8, 35].

In practice, it would be more realistic to minimize *J*_1_ than *J*_2_ since, to our knowledge, personalized cancer vaccines have not shown potential risks for safety or toxicity due to high T cell activation [27, 36, 37]. However, with *J*_2_, we explored the case when minimizing tumor and immune responses simultaneously can lead us to find vaccine doses that fit these two outcomes. In this hypothetical scenario, the excessive T cell response from the cancer treatment could have negative consequences in the context of autoimmune diseases and tissue damage [31, 32]. In autoimmune diseases, the immune system can target and attack healthy cells and tissues due to an overactive T cell response. Moreover, the constant activation and recruitment of T cells can perpetuate the autoimmune process which can cause further damage to affected tissues, leading to impaired organ function, pain, disability, and reduced quality of life in patients. Therefore, regulating and balancing T cell responses during treatment is essential for preventing and managing the adverse effects associated with excessive T cell activation in autoimmune diseases and limiting tissue damage.

Our study has several limitations. First, there are some limitations from the MRM model. The model did not consider potential elimination of tumor cells by activated *CD*4^+^, the functions of memory and regulatory T cells, and tumor eradication by antigen specific antibodies. Moreover, our optimization framework does not allow for optimization of the vaccination schedule. However, we observed that an optimal vaccine dose is usually a combination of higher and lower vaccine doses at some of the scheduled vaccination days. This could open a window of opportunities to explore different schedules where vaccination days requiring low doses (e.g., 0.1 of clinical trial dose) may be removed. Another limitation of our optimization problem is that it does not account for other combined treatments received by the patients. Patients 2 and 6 achieved a positive clinical outcome after receiving anti-programmed cell death protein 1 (anti-PD-1) antibody treatment post vaccination [38]. In the future, it will be important to expand the model such that it accounts for combination therapy and longer outcome [36, 39, 40]. A significant limitation to applying our model in the clinical setting is that the optimization problem relies on the patient longitudinal data over the treatment. Thus, it does not have a predictive value to help determine the vaccine dose prior to the treatment.

Determining the optimal personalized dose of a cancer vaccine is not a straightforward task. There are several logistical complexities involved in vaccine development that may render it impractical for manufacturers to implement the optimal dose. For instance, it may not be feasible to produce the exact optimal number of peptide molecules suggested by the model. The model’s fitting may be inaccurate due to insufficient data to estimate parameters and overfitting, resulting in unreliable and biased outcomes. However, our results offer a promising solution. One can target a cancer vaccine dose to be as effective as the optimal vaccine dose. In addition, our work has the potential to be integrated with a clinical trial, where the optimization framework presented here can be used to”learn” from the outcomes after the initial vaccine doses, and model parameters can be updated continuously over time to make predictions more accurate, like a feedback loop in the digital twin paradigm [41, 42].

This study provides a potential pathway for investigating various dosing regimens in personalized immunotherapy and underscores the significance of comprehending the impacts of alternative dosages in accomplishing the primary objectives of immunotherapy, namely, triggering a potent immune response to reduce or eliminate tumors. Our findings indicate that determining the optimal vaccine dose based on safety and efficacy could assist healthcare practitioners in developing and utilizing cancer vaccine therapies more effectively for each patient. The targeting optimal vaccine doses demonstrated by this paper could serve as a valuable tool for personalized cancer vaccine treatment in a clinical trial setting.

## Acknowledgement

This project was supported in part by an appointment to the Research Participation Program at the U.S. Food and Drug Administration administered by the Oak Ridge Institute for Science and Education through an interagency agreement between the U.S. Department of Energy and the U.S. Food and Drug Administration.

## Disclaimer

This article reflects the views of the authors and should not be construed to represent FDA’s views or policies. S1 Appendix

## Supporting information

## S1 Appendix Model description, necessary conditions and tables of numerical results

